# To Peck or Not To Peck: The influence of early-life social environment on response inhibition and impulsive aggression in Japanese quails

**DOI:** 10.1101/2024.12.20.629368

**Authors:** Alizée Vernouillet, Kathryn Willcox, Reinoud Allaert, Anneleen Dewulf, Wen Zhang, Camille A. Troisi, Sophia Knoch, An Martel, Luc Lens, Frederick Verbruggen

**Author notes:** Correspondence concerning this article should be addressed to Alizée Vernouillet.

## Abstract

Deficits in response inhibition (i.e., the ability to suppress inappropriate responses) may contribute to increased impulsive aggression (i.e., unplanned behaviors that harm others). Since early-life environment might influence the development of response inhibition, it could also indirectly affect impulsive aggression. However, this relationship has rarely been directly examined. Here, we investigated whether response inhibition is associated with impulsive aggression and whether this relationship explains the effects of early-life social environments on impulsive aggression in juvenile Japanese quails (*Coturnix japonica*). Quails (*n* = 120) were raised in two social conditions: Small groups of five birds or Large groups of 15 birds. Response inhibition was assessed using the Barrier and Cylinder tasks, while impulsive aggression was measured in two contexts - within a group of five familiar individuals and in a dyad with an unfamiliar individual. We found that some aspects of response inhibition were related to impulsive aggression. Furthermore, quails reared in small groups showed significantly poorer response inhibition than those reared in large groups. Yet, impulsive aggression did not significantly differ between the two conditions. These findings suggest that, while impulsive aggression is partly related to response inhibition, other factors mitigate the effects of early social environments on its expression.

## 2. Introduction

Group living brings many benefits, such as reducing predation pressure and increasing foraging efficiency, but can also create conflicts between members due to increased competition (1). These agonistic interactions between individuals may result in *impulsive aggression* (i.e., the unplanned behavioural response of inflicting damage to self or others, (2)), that in turn may sometimes have negative fitness consequences for one or multiple individuals involved (e.g., reduced access to resources, injuries, or even death). While a lot of research has focused on aggression as a personality trait in animals (as reviewed by e.g., (3)), little is known about the cognitive mechanisms underlying aggression.

Some researchers suggest that impulsive aggression results from deficits in *response inhibition* (also referred to in the animal cognition field as *inhibitory control*; (4–6); but see (7)), the suppression of inappropriate pre-potent behavioural responses or actions (8–10). Indeed, inhibition deficits are correlated with increased aggression in rats (11), dogs (12), long-tailed macaques (13), and humans (14–16). Additionally, both increased levels of aggression and decreased levels of response inhibition have been correlated with lower levels of central serotonin (5-HT), a well-conserved neuromodulator of nervous systems in both invertebrates and vertebrates (9,11,17,18). However, most studies investigating the link between response inhibition and impulsive aggression are correlational. This has two important consequences. First, response inhibition deficits could cause or contribute to impulsive aggression, but it is also possible that a common underlying factor simply influences both independently (as suggested by (19)). Second, it has been proposed that environmental factors during early life contribute to response inhibition deficits and impulsive aggression later in life. Yet again, causal relationships have been hard to establish, and systematic research is lacking.

Early-life environmental conditions, both physical and social, can shape an individual’s cognitive abilities and personality (e.g., (20–24)). For example, response inhibition is key to maintain spatial and temporal cohesion between group members (25). Indeed, individuals within a group benefit from being surrounded by others and performing the same activity together, so they need to adjust their activity to that of other group members (e.g., (26,27)). Additionally, within a group, subordinate individuals might inhibit feeding or mating in the presence of dominant individuals to avoid potential aggressive interactions or confrontations (28). Given that social complexity is multi-faceted and highly variable, both within and between species (29), in the present study we focused on one key aspect of the social organization – social group size (30). An increase in group size means an increase in potential interactions with other group members, and therefore an increase in possibilities to learn how to interact with other individuals. Social group size has been shown to correlate with higher levels of response inhibition in Australian magpies (*Cracticus tibicen dorsalis*, (31)), in hyenas (*Crocuta crocuta*, (32)), and in primates (33), but not in guppies (*Poecilia reticulata*, (24,34)). Hence, we expected that a larger group size required individuals to develop better response inhibition due to greater social requirements.

Many studies have found aggression is usually suppressed in larger groups (e.g., calves – (35), chickens – (36,37), fish – (38), pigs – (39)). Based on the same reasoning described in the previous paragraph, with larger groups come a higher number of potential competitors for resources, and individuals might thus benefit to switch to a more socially tolerant strategy (e.g., (37,40), but see e.g., (41)). Yet, methodological differences between studies make it difficult to determine whether, and if so in what direction, early-life social environment shapes an individual’s aggressive tendencies (42). Based on previous research and on the theoretical link between inhibition and aggression, and because we predicted that individuals raised in larger groups would have better response inhibition, we expected levels of impulsive aggression to decrease with increasing group size.

In this study, we investigated the link(s) between early-life social environment, response inhibition and aggression in juvenile Japanese quails (*Coturnix japonica*). Japanese quails are a precocial social bird species that is easy to raise in captivity, which enabled us to rear them without the presence of adult quails. This controlled raising environment allowed us to manipulate their postnatal social conditions, while controlling all other environmental parameters. Additionally, Japanese quails have long been recognized as an ideal model for studying aggression, as hierarchies within their social group are established and maintained through frequent aggressive interactions (e.g., (26,43,44)). As a result, their typical behaviors, including aggressive behaviors towards conspecifics, have been extensively documented (e.g., (44–46)). These combined factors were crucial in establishing causal relationships between early-life social environment and an individual’s response inhibition and aggression.

We evaluated response inhibition in Japanese quails with two commonly used tasks to evaluate an animal’s response inhibition: the *Barrier task*, during which individuals must detour around a vertical transparent barrier to reach a reward, and the *Cylinder task*, during which individuals must reach through the openings of a transparent cylinder to obtain a reward. Both tasks are purporting to measure the same aspect of response inhibition and have been referred to as “detouring” tasks or “motoric self-regulation” tasks, as, in both tasks individuals must inhibit a prepotent motor action (i.e., going towards the visible reward) and instead change it into a different one (i.e., detouring around a physical barrier) (47) (but see (48)). Aggression was evaluated in two different contexts: once in a group of familiar individuals (e.g., (44)), and once in presence of an unfamiliar individual (e.g., (49)).

The main goal of the study was to experimentally determine the influence of early-life social environment on response inhibition and impulsive aggression in Japanese quails. We focused on three research questions: (A) do deficits in response inhibition cause impulsive aggression at an individual-level, and (B) does social group size during early life shape an individual’s response inhibition and (C) impulsive aggression? Based on previous correlational research and on theoretical models, we predicted that, if indeed higher impulsive aggression can be attributed to a deficit in response inhibition, (A) response inhibition and impulsive aggression would be inversely related at an individual-level, and that (B and C) individuals raised in larger groups would have a higher level of response inhibition and be less aggressive than individuals raised in smaller groups.

## 3. Material and methods

### Subjects

We used 120 juvenile Japanese quails (*n* = 60 females, *n* = 60 males) in this study. Birds were hatched simultaneously over two days and kept together at the Opvangcentrum Vogels en Wilde Dieren, Ostend, Belgium. Birds were housed in four identical heated lofts, each loft containing four 1m^2^ (1 m length x 1 m width x 2 m height) enclosures. Each enclosure included an electric hen heating plate (25 cm length x 25 cm width, Comfort Chicks), which was replaced with a plastic cover when birds were 16 days old. Outside of the experimental food restriction procedure, birds were provided with *ad lib* grain food mix (Aveve), presented in a single 50-cm long linear chick feeder (Voss Framing). Birds had free access to water. Each loft followed a 12:12 day-night cycle, with light onset at 07:00 am. All birds were visually checked at least twice a day (once in the morning and once in the evening). Measurements (weight and tarsus length) were taken when the birds were 20 days old (before the start of the study) and at 44 days old (at the end of the study). Quail care was performed by AV, KW, RA, AD, WZ, and CT, under AV and RA’s supervision.

Within each loft, one of the four enclosures housed a large group of 15 individuals (Large), and the three other enclosures housed a small group of five individuals (Small). On day 2 post-hatch, birds were randomly assigned to either a Small or a Large enclosure. Hence, 60 birds were assigned to each experimental condition. Previous research has found differences in social bonding and other social affiliation behaviors between individuals raised in groups of 6 vs 15 (50). Additionally, wild Japanese quails have broods of up to 15 individuals (51,52), and once sexually mature, they can be monogamous or live in small flocks (51). Hence, we predicted that Japanese quails raised in groups of five and 15 would differ in their RI and IA if the early-life social group size influenced the development of cognitive and behavioral traits.

### General Experimental Procedures

Quails were habituated to being handled and to the testing environment during their second and third week. Testing started when birds were 24 days old and ended when they were 41 days old. At that age, juvenile quails are independent and can be socially isolated without showing signs of distress. All 120 individuals underwent the Barrier task, the Cylinder task, the Familiar Group task, and the Unfamiliar Individual task, given in that order for all individuals. Birds were tested individually during the Detour and the Cylinder tasks (i.e., response inhibition tasks) at the age of 24-33 days old, and with other individuals during the Familiar Group and the Unfamiliar Individual tasks (i.e., impulsive aggression tasks) at the age of 39-41 days old. Tasks were performed on separate days. A schematic overview of the study timeline can be found in the Supplementary Material (Fig. S1).

Testing took place between 08:30 and 14:00 (local time). Food was removed the evening prior testing to ensure motivation to participate in the experiment. On the morning of testing, birds were individually placed in a portable cage with multiple compartments (Ducatillon) and transported to a waiting room, where they were kept before and after testing until all individuals had been tested.

All testing occurred in a test box (86 cm length x 146 cm width x 91 cm height). The testing box was lit with LED lights to control for lighting. All trials were recorded using a camera (BASCOM) mounted to the ceiling of the test box. A ‘start box’ (29 cm length x 45 cm width x 29 cm height) was attached to the front of the test box. The start box had two doors – one transparent and one opaque – which slid vertically on rails between the two boxes, and a separate light. Inside the start box was another square piece of wood that served as the back of the box attached to a handle, which could slide forward and was used to gently push the birds into the test box when required.

Every testing trial followed the same general procedure. First, the bird(s) to be tested was (were) placed in the start box with both start box doors closed. Next, the opaque door was opened, allowing the bird(s) to see within the test box but not enter the test box. After 15 seconds the transparent door was opened, indicating the start of the trial. Birds that did not leave the start box within 30 seconds were gently pushed into the test box. Once the bird(s) either reached the reward location, or the time limit for the trial was reached, the test box lights were turned off, indicating the end of the trial. The bird(s) was (were) given 15 seconds to return to the start box; if it(they) did not, it was (they were) caught by the experimenter and placed back in the start box. Once all birds completed all required trials for a session, they were returned to their home enclosures. Data collection was performed by AV, KW, RA, AD, WZ, CT, and LL, under AV and KW’s supervision. Videos were coded using BORIS v.7.13 (Behavioral Observation Research Interactive Software (53)). All the videos of the response inhibition tasks were coded by one experimenter (KW), who was naïve to the conditions. All the videos of the impulsive aggression tasks were coded by another coder (SK), who was naïve to the conditions and the experimental hypotheses. To assess inter-observer reliability, 20 % of the videos of the response inhibition tasks were coded by one coder (SK), who was naïve to the conditions and the experimental hypotheses, and 20 % of the videos of the impulsive aggression tasks were coded by one experimenter (AV). Inter-rater reliability was calculated using BORIS, and was high, with average Cohen’s kappa coefficients of 0.840 and 0.805, for all response inhibition videos and all impulsive aggression videos respectively (54).

### Habituation

Birds were gradually habituated to the portable cage from 20-days-old onwards and to the test box at 22-23 days old. For the habituation in the test box, a feeder was placed 50cm from the test box entrance. The feeder used was filled with grains and was the same design as those used in the birds’ enclosures, but smaller (30cm length). A petri dish with two live mealworms was also placed in front of the feeder. Both rewards were familiar to the birds and were used to entice them to participate in testing. The total trial time was 5 minutes; each bird received one habituation trial.

### Response inhibition tasks

#### Barrier task

*Apparatus*. The apparatus for the Barrier task consisted of a square wooden frame (40 cm length x 41.2 cm height) with two horizontal supports (20 cm width) (Fig. 1). The frame was placed 55 cm from the entrance of the test box, centrally such that there was 22 cm between it and the test box wall on each side. A reward feeder filled with grains was placed directly behind the barrier, and a petri dish containing two mealworms was placed directly behind the feeder. An opaque barrier was used during the training trials and a transparent barrier was used during the testing trials, so the reward food was visible to the birds through the barrier during the test session (Fig. 1). Both barriers had a strip of green tape on the left side and a strip of yellow tape on the right side (the same colours as the enclosure and reward feeders).

**Figure 1.**
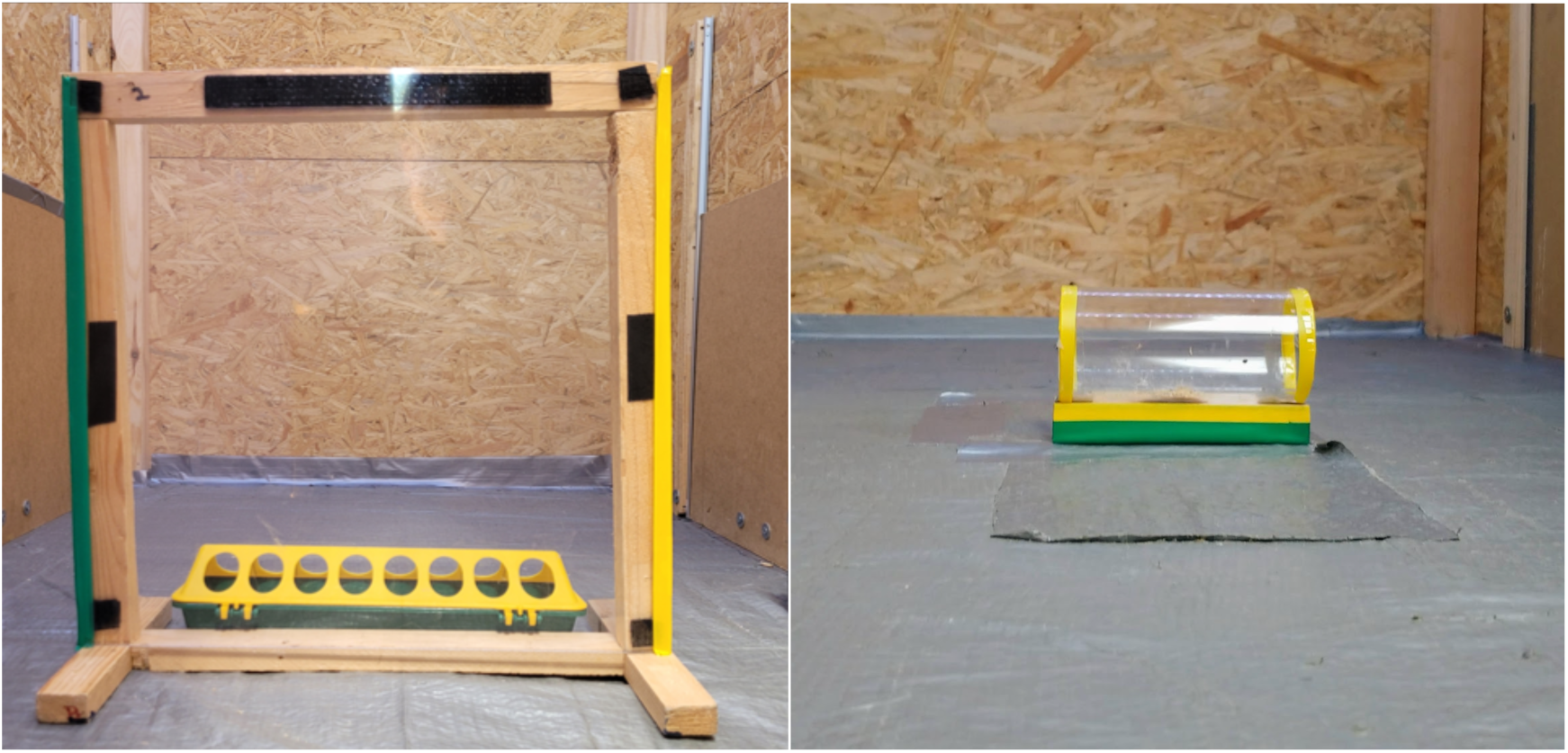
Apparatuses used for the *Barrier task* (left) and for the *Cylinder task* (right) with Japanese quails (*n* = 120).

*Procedures*. This task included a training session and a testing session on two separate, consecutive days.

During the training session, the opaque barrier was attached to the frame. During the first trial, the bird was given two minutes to detour around the barrier to reach the food; from the second trial onwards, it had one minute and a half. Trials were repeated until the bird successfully detoured on three consecutive trials, with a maximum of seven trials per bird. If the bird did not detour once within three consecutive trials, the session ended. All birds passed the training session.

The testing session on the following day consisted of one training trial and one test trial per bird. The test trial followed the same procedure as during the training trials, but the transparent barrier was used instead of the opaque barrier.

> ‘Successful’ trials were defined as when the bird detoured around the barrier (opaque or transparent) without interacting (i.e., pecking or pushing) with it.

#### Cylinder task

*Apparatus*. The cylinder apparatus consisted of a transparent Plexiglas tube with open ends (5.4 cm diameter x 13.2 cm length) mounted on a small wooden platform (13 cm length x 3 cm width x 2 cm height; Fig. 1). The wooden base was covered with horizontal strips of yellow and green tape. The apparatus was placed 40 cm from the entrance of the test box, so that the entrances were perpendicular to the start box and secured to the floor with tape. One freshly killed mealworm and 1tsp of grain were placed centrally inside the cylinder. Quails had to put their head within the cylinder to retrieve the food.

*Procedures*. The Cylinder task occurred during a single test day. During each trial, the bird had two minutes to reach the food inside the cylinder. Trials were repeated until the bird completed ten trials, which was the maximum number of trials given, or until the bird failed to reach the food for three consecutive trials.

‘Successful’ trials were defined as when the bird reached the food through one of the two open sides of the cylinder without first pecking at the outside of the cylinder. For the purpose of this study, we only focused on the first trial, as we believe it best measure the stopping behaviour of an individual (55). Indeed, the subsequent trials might reflect the inhibitory learning of the individual and is investigated in a different study (56).

#### Behavioural measures of response inhibition tasks

For each quail, the dependent measures evaluated were (1) trial success (i.e., whether the trial was a success), and (2) time spent ‘interacting’ with the barrier/cylinder (i.e., time the bird spends actively and repeatedly pecking/pushing the barrier).

### Impulsive aggression tasks

Impulsive aggression tasks had multiple trials and different experimental phases, which allowed us to investigate whether quails’ aggressive behaviour was repeatable across contexts differing in food availability. The same set of birds were tested for all trials. This repeatability aspect is being investigated in a different project and not analysed in this study (Vernouillet et al. in prep.). Here, we focused on the behaviour of individuals during the first trial.

#### Familiar Group

*Apparatus*. A feeder, identical to the one present in the home enclosure of individuals and filled with grains, was placed in the middle of the text box. A sheet of semi-transparent plastic was placed inside the feeder to control access to the food. The feeder was present through the duration of the trial, but access to the food was adjusted during the trial (see Procedures).

*Procedures*. Five quails from the same enclosure were tested together for two trials on two consecutive days. The five individuals were placed in the start box and, following the same procedure as during the previous tasks, were let into the test box after 15 seconds after the opaque door of the start box was removed. Trials lasted 10 minutes. During the first trial, access to the feeder was completely thwarted for the first five minutes of the trial (No Food phase), with the sheet of plastic blocking access to all the holes in the feeder, then partially thwarted so that only one individual could access the food for the last five minutes of the trial (Limited Access phase), with the sheet of plastic blocking access to all but one hole in the feeder. During the second trial, the procedure was reversed (i.e., Limited Access phase during the first five minutes, No Food phase during the last five minutes).

#### Unfamiliar Individual

*Apparatus*. Similar to the Familiar Group task, a feeder, identical to the one present in the home enclosure of individuals and filled with grains, was placed in the middle of the test box. A sheet of semi-transparent plastic was placed inside the feeder to control access to the food. The feeder was placed inside the test box during the trial (see Procedures).

*Procedures*. Two quails from a different enclosure and loft (but of the same experimental condition) were tested together for three trials on three consecutive days. Two individuals in the start box, and following the same procedure as during the precedent tasks, were let into the test box after 15 seconds from removing the opaque door of the start box. Trials lasted three minutes. The feeder was placed in the test box after one minute. During the first trial, only one bird had access to food when the feeder was placed in the test box, with the sheet of plastic blocking all but one hole in the feeder. During the second trial, both birds could freely access the food when the feeder was placed in the test box, as there was no sheet of plastic blocking access to the feeder. During the third trial, none of the birds had access to the food, as access was thwarted, with the sheet of plastic blocking all holes in the feeder.

#### Behavioural measures of impulsive aggression tasks

For each quail, the dependent measures evaluated were (1) number of pecks given, (2) number of pecks received, (3) time spent chasing, (4) time spent escaping, and (5) time spent pushing. Descriptions of each behaviour can be found in Table S1.

### Statistical analyses

We first wanted to perform a structural equation modelling approach (SEM) to determine whether response inhibition, as estimated by the measures assessed during the Barrier and the Cylinder tasks, could predict impulsive aggression, as estimated by the behavioral measures assessed during the Familiar Group and the Unfamiliar Individual tasks, and whether early-life social condition influenced response inhibition and impulsive aggression. However, a preliminary Confirmatory Factor Analysis indicated a lack of correlations among measures assessing the latent factors of interest (i.e., response inhibition and impulsive aggression), and none of the models tested had a decent fit (see Supplementary Material, Figures S2 and S3, Table S2 for more details on factors analyses and SEM).

We thus used a different approach for our analyses. First, to determine whether both the Barrier task and the Cylinder task were assessing the same aspect of response inhibition, we correlated (1) first trial success using a tetrachoric correlation (for binary data), and (2) time spent ‘interacting’ with the barrier/cylinder using a Pearson’s correlation. Second, to determine whether both impulsive aggression tasks were assessing the same aspect of aggression, we conducted a PCA with (1) number of pecks given, (2) number of pecks received, (3) time spent chasing, (4) time spent escaping, and (5) time spent pushing for both tasks. PCA was based on a correlation matrix. We only considered components whose Eigenvalues were higher than 1 and for each component, we only considered loadings whose contribution was higher than what was expected if all loadings were of equal weight. Third, as we did not find any significant correlation between the different response inhibition variables (see Results), we assessed whether higher levels of impulsive aggression were linked to response inhibition deficits by evaluating whether aggression scores (as calculated during the PCA) significantly differ between individuals that were successful during the first trial of each response inhibition task using a Student t-test and whether they correlated with time spent ‘interacting’ with the barrier/cylinder using a Pearson’s correlation. Importantly, unlike the SEM approach, this approach does not allow us to infer the direction of the potential link between response inhibition and impulsive aggression.

To determine whether early-life social environment influenced an individual’s response inhibition and/or impulsive aggression, we used a generalized linear model (GLM) approach, with (1) trial success (i.e., whether the barrier/cylinder testing trial was successful) for each response inhibition task separately, and (2) time spent ‘interacting’ with the barrier/cylinder (i.e., time the bird spends actively and repeatedly pecking/pushing the barrier) for each response inhibition task separately, as dependent variables associated with response inhibition, and (3) using the PCA scores obtained for the first two axes, as aggressive behaviours during the Unfamiliar Individual task mostly loaded on PC1 and aggressive behaviours during the Familiar Group task mostly loaded on PC2 (see Results) as dependent variables associated with impulsive aggression. Full models included social group condition, as well as morphological factors such as body weight and sex.

To assess whether specific factors influenced our dependent variables, we compared full models to a reduced model that did not include the factor that was evaluated (following the procedures described by (57,58)). Parameter estimation was achieved using the residual maximum likelihood ratio test.

For trial success during the Barrier and the Cylinder tasks, we used a binomial distribution. For time spent interacting with the barrier/cylinder, we rounded the values to the nearest second and used a Poisson distribution. We first checked the assumptions of the GLMs were met using the *DHARMa* package (59). Models were checked for dispersion and zero-inflation using residual plots. In case residuals were over-dispersed, we added a dispersion parameter (this was the case for the GLM on the second impulsive aggression score). In case there were too many zero in our dataset, we added a zero-inflated model with a constant at the origin, and in the case that the assumptions of the models were still not met, we changed the family to a negative binomial family (this was the case for the GLM on time spent pecking for both response inhibition tasks).

Analyses were conducted using R version 4.3.2 (60) with the *glmmTMB* (61), *psych* (62), *FactoMineR* (63), and *factoextra* (64). Descriptive statistics were obtained using the *Rmisc* package (65). Plots were created using the *ggplot2* (66), *ggsignif* (67), *ggpubr* (68), *patchwork* (69), *wesanderson* (70) packages. Alpha was set at 0.05 for all statistical analyses. All statistical analyses were performed by AV, verified by KW and CAT.

## 4. Results

### Overview of response inhibition tasks

#### Overview of Barrier task

During the testing trial, 19 individuals detoured successfully (i.e., without touching the transparent barrier), 93 individuals detoured unsuccessfully (i.e., after touching the transparent barrier), seven individuals touched the barrier but did not detour during the entire duration of the trial, and one did not interact with the apparatus nor detoured around the transparent barrier (view Table S3 for an overview of the response outcomes of the quails based on their social group and on sex). Among individuals who pecked the barrier (*n* = 100), average time spent pecking the barrier was 17.5 (±*SE*: 1.4) seconds.

#### Overview of Cylinder task

During the first testing trial, 14 individuals detoured successfully (i.e., without touching the transparent cylinder), 46 individuals detoured unsuccessfully (i.e., after touching the transparent cylinder), 36 individuals touched the cylinder but did not detour, and 24 did not interact with nor detoured around the cylinder (view Table S3 for an overview of the response outcomes of the quails based on their social group and on sex). Among individuals who pecked the cylinder (*n* = 82), average time spent pecking the cylinder was 15.0 (±*SE*: 1.6) seconds.

#### Repeatability of response inhibition

There were 88 individuals who were not successful during either response inhibition task, 18 individuals who were successful during the Barrier task only, 13 individuals who were successful during the Cylinder task only, and only one individual who was successful during both response inhibition tasks.

For this analysis, only individuals who interacted (either by detouring around and/or by pecking the transparent barrier) during both tasks were included (*n* = 95). Results from the correlations indicated that both tasks were measuring different aspects of RI, as neither success between tasks (tetrachoric correlation coefficient: -0.250) nor time spent interacting with the transparent barrier/cylinder (Pearson correlation coefficient: r = -0.116, *t* = -1.129, *df* = 93, *p* = 0.262, Fig. 2) correlated between tasks.

**Figure 2.**
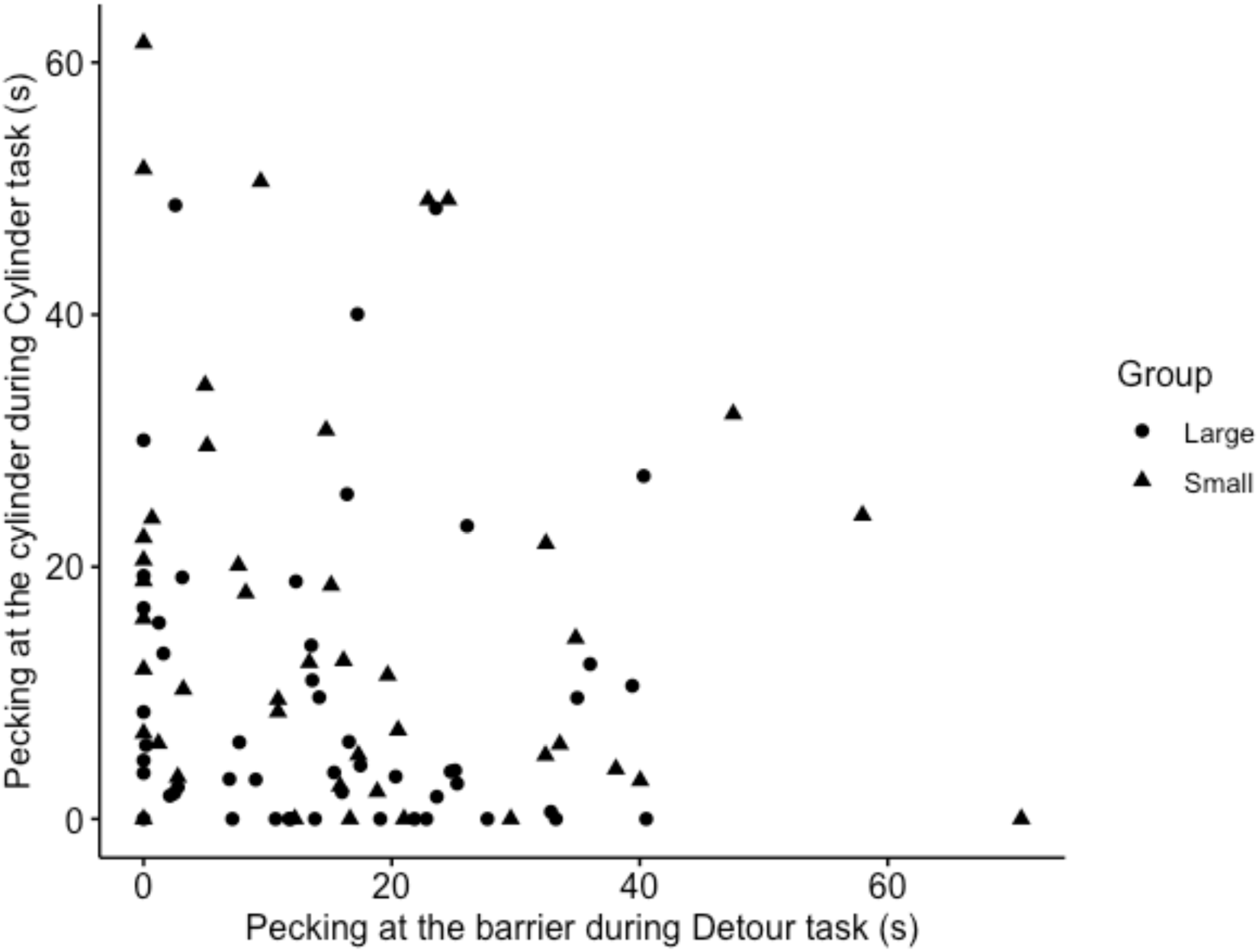
Time spent pecking at the transparent barrier during the Barrier task did not significantly correlate with the time spent pecking the cylinder during the Cylinder task in Japanese quails (Pearson correlation coefficient: r = -0.116, *t* = -1.129, *df* = 93, *p* = 0.262).

### Overview of impulsive aggression tasks

#### Overview of Familiar Group task

All 120 individuals participated during the Familiar Group task. Quails gave on average 8.2 ± 16.7 pecks, received on average 8.2 ± 9.0 pecks, chased another individual for 1.0 ± 5.0 seconds, escaped for 1.7 ± 4.4 seconds, and pushed others for 2.3 ± 6.0 seconds.

#### Overview of Unfamiliar Individual task

All 120 individuals participated during the Unfamiliar Individual task. Quails gave on average 2.5 ± 5.3 pecks, received on average 2.5 ± 5.3 pecks, chased another individual for 1.8 ± 5.7 seconds, escaped for 3.3 ± 7.5 seconds, and pushed the other for 0.3 ± 0.9 seconds.

#### Repeatability of impulsive aggression

All individuals were included in the analysis, as they all participated in both tasks (*n* = 120). Results of the PCA indicated that there was not a unique impulsive aggression aspect measured during the tasks (Table 1). The first five components had an Eigen value higher than 1 and together explained 78.1 % of the variance in the data. We found that both tasks investigated a different impulsive aggression aspect, as measures related to each task loaded on different components (Table 1). For the subsequent analyses, we used the individual scores obtained on the first two components, as they best represent the aggression scores during each task (PC1 for Unfamiliar Individual, PC2 for Familiar Group).

**Table 1.**
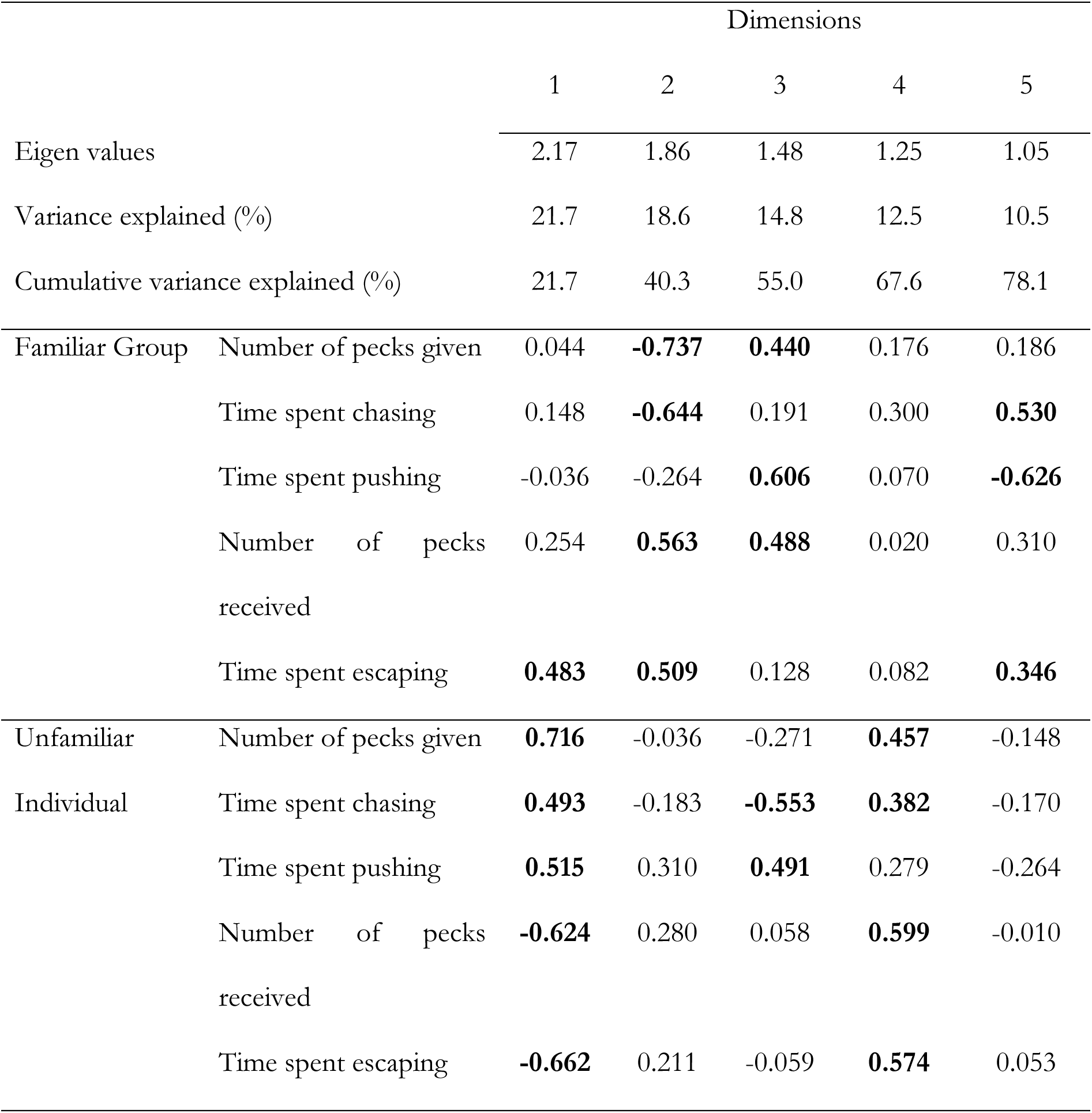
PCA loadings for the measures on impulsive aggression in Japanese quails. Loadings considered important (i.e., whose contribution was higher than what was expected if all loadings were of equal weight) are indicated in bold.

### Relationship between response inhibition and impulsive aggression

We evaluated whether the different response inhibition behavioural measures were related to the aggression scores of the individuals. For these analyses, we used reduced datasets that included quails that interacted with the apparatuses (either by detouring successfully or by pecking at the apparatus) either during Barrier task (*n* = 119) for measures associated to that task or during the Cylinder task (*n* = 96) for measures associated to that task.

Aggression scores for the Unfamiliar Individual task (PC1) significantly correlated with the amount of time spent pecking on the cylinder during the Cylinder task. Individuals that spent more time pecking at the cylinder had a higher aggression score than individuals that spent less time pecking at the cylinder (Pearson coefficient correlation: *r* = 0.221, *t* = 2.199, *d.f.* = 94, *p* = 0.030; Fig. 3). Unsuccessful individuals tended to have higher aggression scores in in comparison to successful individuals (*t* = 2.072, *d.f.* = 18.08, *p* = 0.053). Response inhibition measures assessed during the Barrier task did not significantly correlate with the PC1 aggression scores for that task (Success during the Barrier task: *t* = -0.090, *d.f.* = 38.55, *p* = 0.929; Time spent pecking the transparent barrier: Pearson coefficient correlation: *r* = -0.018, *t* = -0.197, *d.f.* = 117, *p* = 0.844).

**Figure 3.**
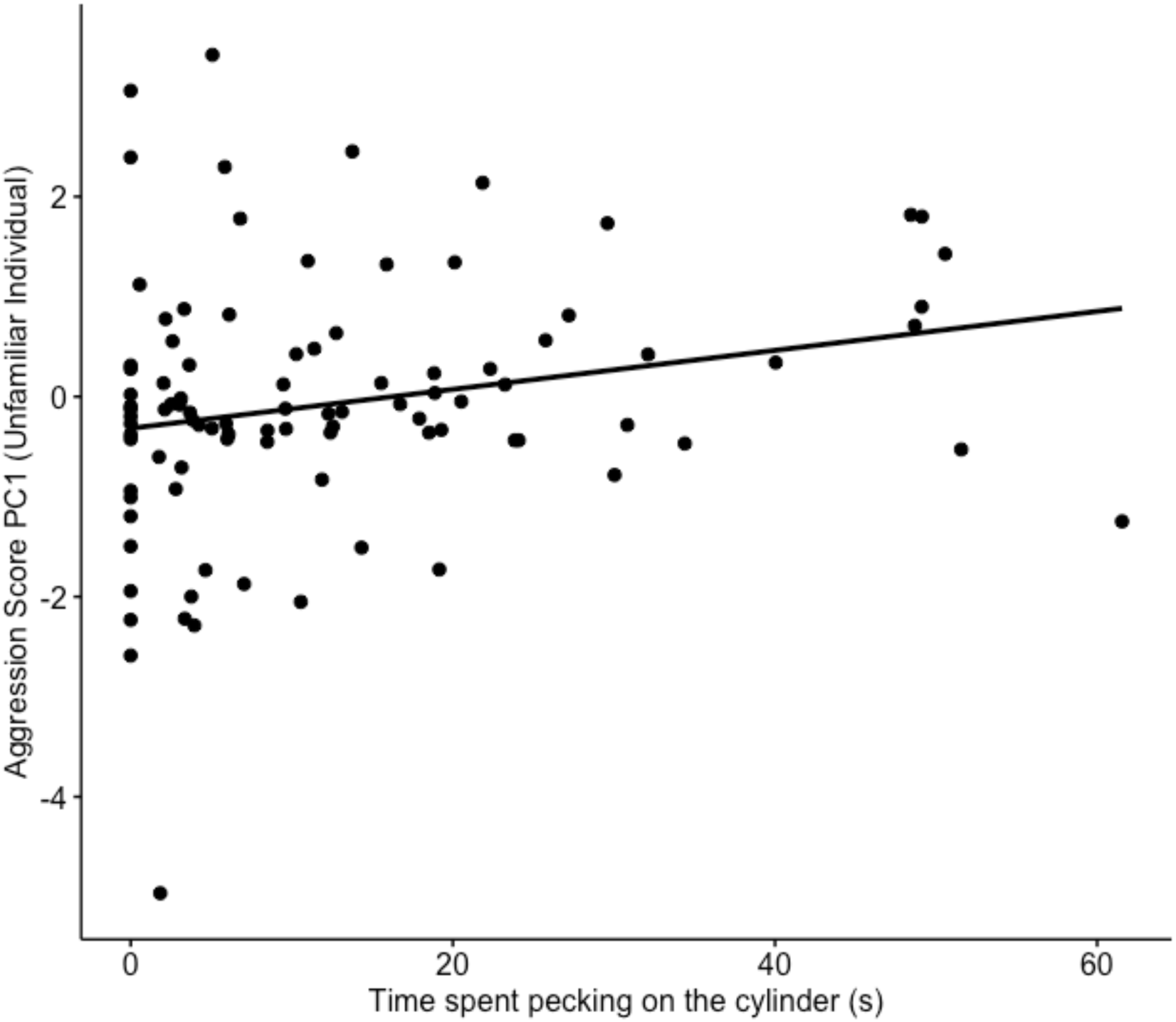
Time spent pecking on the cylinder during the Cylinder task significantly predicted the aggression scores obtained during the Unfamiliar Individual task (Pearson coefficient correlation: *r* = 0.221, *t* = 2.199, *d.f.* = 94, *p* = 0.030).

Aggression scores for the Familiar Group task (PC2) were significantly linked to the success during the Barrier task, with individuals that successfully detoured during the Barrier task being more aggressive than individuals that were unsuccessful (*t* = 2.757, *d.f.* = 20.79, *p* = 0.012; Fig. 4). No other response inhibition measures significantly influenced the PC2 aggression scores (Success during the Cylinder task: *t* = 1.588, *d.f.* = 20.54, *p* = 0.128; Time spent pecking the transparent barrier: *t* = -1.757, *d.f.* = 117, *p* = 0.082; Time spent pecking the transparent cylinder: *t* = -1.645, *d.f.* = 94, *p* = 0.103).

**Figure 4.**
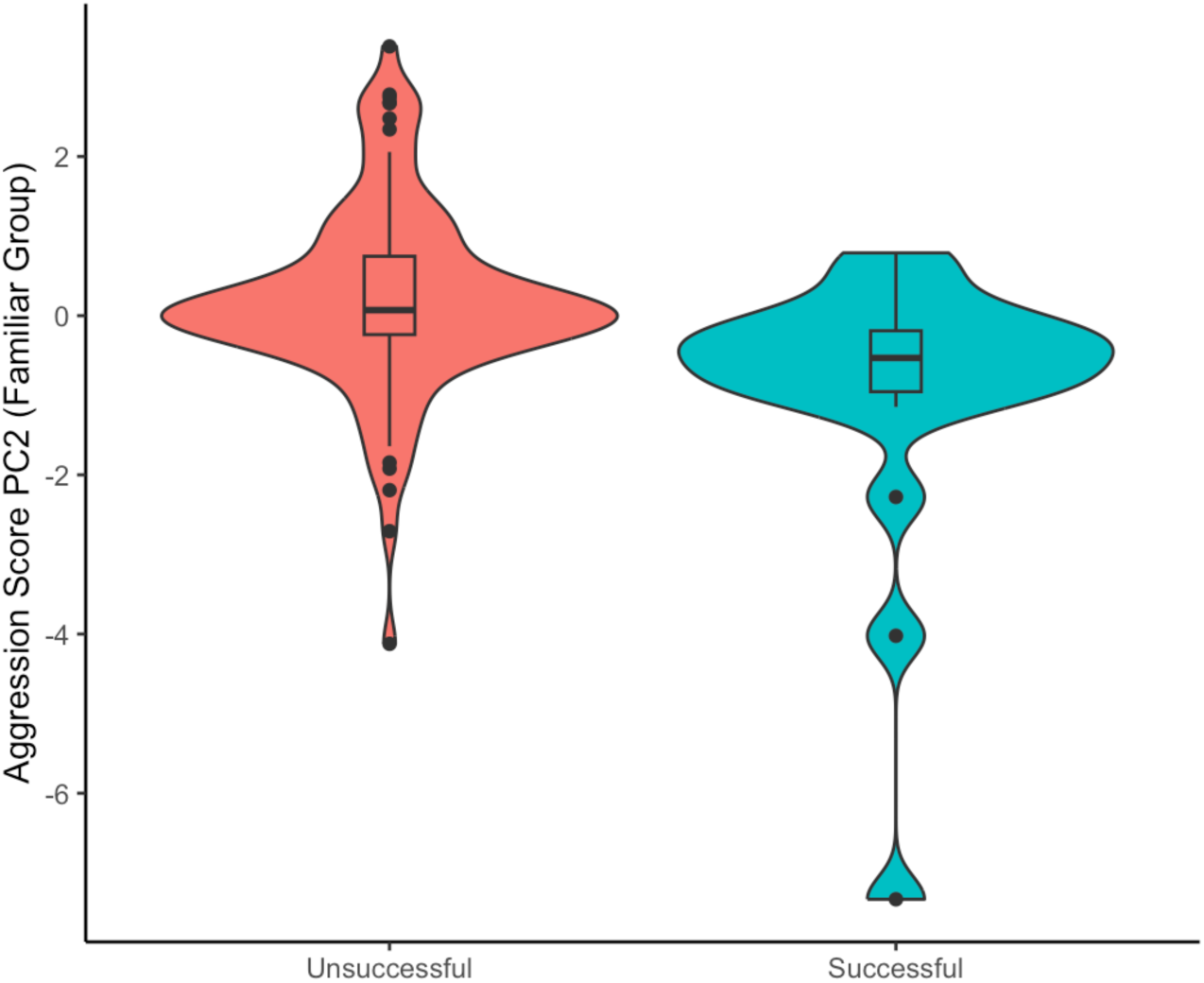
Success during the Barrier task significantly predicted the aggression scores PC2 obtained during the Familiar Group task. Note that for this aggression score, negative scores indicate more aggressive behaviours than positive scores.

### Influence of group size on response inhibition and impulsive aggression

#### Influence of group size on response inhibition

Given the lack of contextual repeatability, we evaluated the effects of Group Size, Sex, and Body Weight on success and on the time spent interacting with the transparent barrier/cylinder for both Response Inhibition tasks separately.

Only individuals who interacted (either by detouring around and/or by pecking the transparent barrier) were included for the analyses on success (*n* = 119 for the Barrier task, *n* = 96 for the Cylinder task; see Table S3 for overview of number). Success during the Barrier task and success during the Cylinder task were not explained by Group Size, Sex, nor Body Weight (Table 2).

**Table 2.** Factors evaluated to explain success (i.e., successfully detouring without touching the barrier or not) and time spent pecking during the Barrier task and during the Cylinder task. Significant factors are indicated in bold.

|  | Barrier task |  |  |  |  |  | Cylinder task |  |  |  |  |  |
| --- | --- | --- | --- | --- | --- | --- | --- | --- | --- | --- | --- | --- |
|  | Success |  |  | Time spent pecking |  |  | Success |  |  | Time spent pecking |  |  |
| | $\chi^2$ | d.f. | p | $\chi^2$ | d.f. | p | $\chi^2$ | d.f. | p | $\chi^2$ | d.f. | p |
| Group Size | 0.732 | 1 | 0.392 | 1.608 | 1 | 0.205 | 0.421 | 1 | 0.517 | **6.256** | **1** | **0.012\*** |
| Sex | 3.725 | 1 | 0.054 | 0.063 | 1 | 0.802 | 1.410 | 1 | 0.235 | 3.719 | 1 | 0.054 |
| Body Weight | 1.298 | 1 | 0.255 | **5.835** | **1** | **0.016\*** | 0.002 | 1 | 0.965 | **14.268** | **1** | **< 0.001\*** |

Only individuals who pecked at the transparent barrier were included for the analyses on time spent pecking (*n* = 100 for the Barrier task, *n* = 82 for the Cylinder task; see Table S3 for overview of number). Time spent pecking at the transparent barrier during the Barrier task was not explained by Group Size (Fig. 5) nor by Sex (Table 2). However, Body Weight significantly influenced time spent pecking at the transparent barrier, with individuals with a lower weight spending more time interacting with the barrier than individuals with a higher weight (Fig. S4). Group Size significantly influenced the time spent pecking at the transparent cylinder during the Cylinder task, with individuals in the Large group spending less time pecking at the transparent cylinder than individuals in the Small group (Time spent pecking at the transparent cylinder *M ± SE*: Large group: 11.64 ± 1.92 s, Small group: 18.70 ± 2.50 s; Table 2; Fig. 5). Body Weight also significantly influenced time spent pecking at the cylinder, with individuals with a higher weight spending more time interacting with the cylinder than individuals with a lower weight (Figure S4). There was no significant difference between males and females (Table 2).

**Figure 5.**
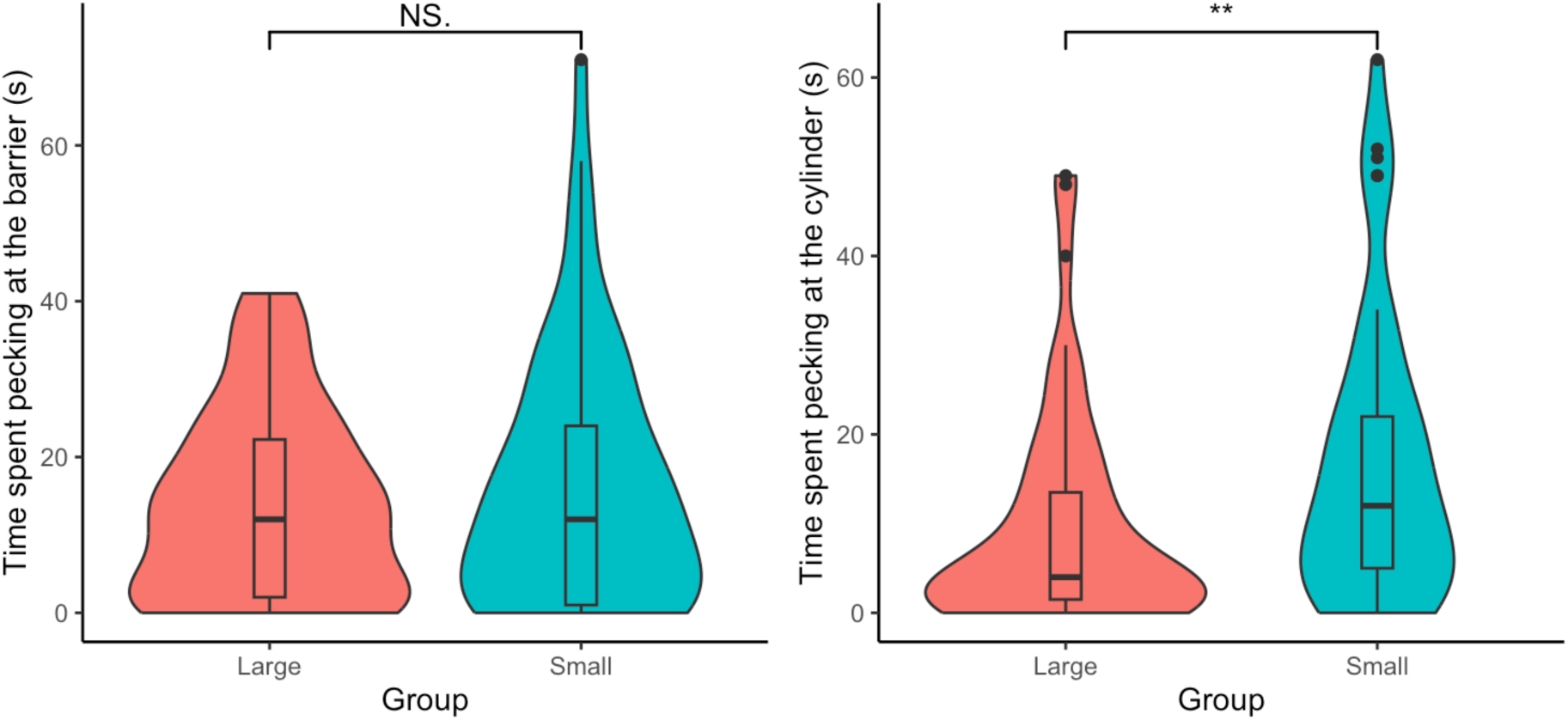
Left. Time spent pecking at the transparent barrier during the Barrier task did not significantly differ between social groups (*χ^2^* = 1.608, *d.f.* = 1, *p* = 0.205). Right. Quails in the Large group spent significantly less time pecking at the transparent cylinder than quails in the Small group (*χ^2^* = 6.256, *df* = 1, *p* = 0.012).

#### Influence of group size on impulsive aggression

Given the lack of repeatability in impulsive aggression between contexts, we evaluated the effects of Group Size, Sex, Body Weight on the aggressive scores for both impulsive aggression tasks separately (*n* = 120 quails).

Aggression scores for the Unfamiliar Individual task (PC1) were not explained by Group Size (*χ^2^* = 0.001, *d.f.* = 1, *p* = 0.973; Fig. 6) nor by Body Weight (*χ^2^* = 1.178, *d.f.* = 1, *p* = 0.278). However, male quails had a significantly higher aggression score than female quails (Females: -0.28 ± 1.54, Males: 0.28 ± 1.38; *χ^2^* = 4.318, *d.f.* = 1, *p* = 0.038; Figure S5).

**Figure 6.**
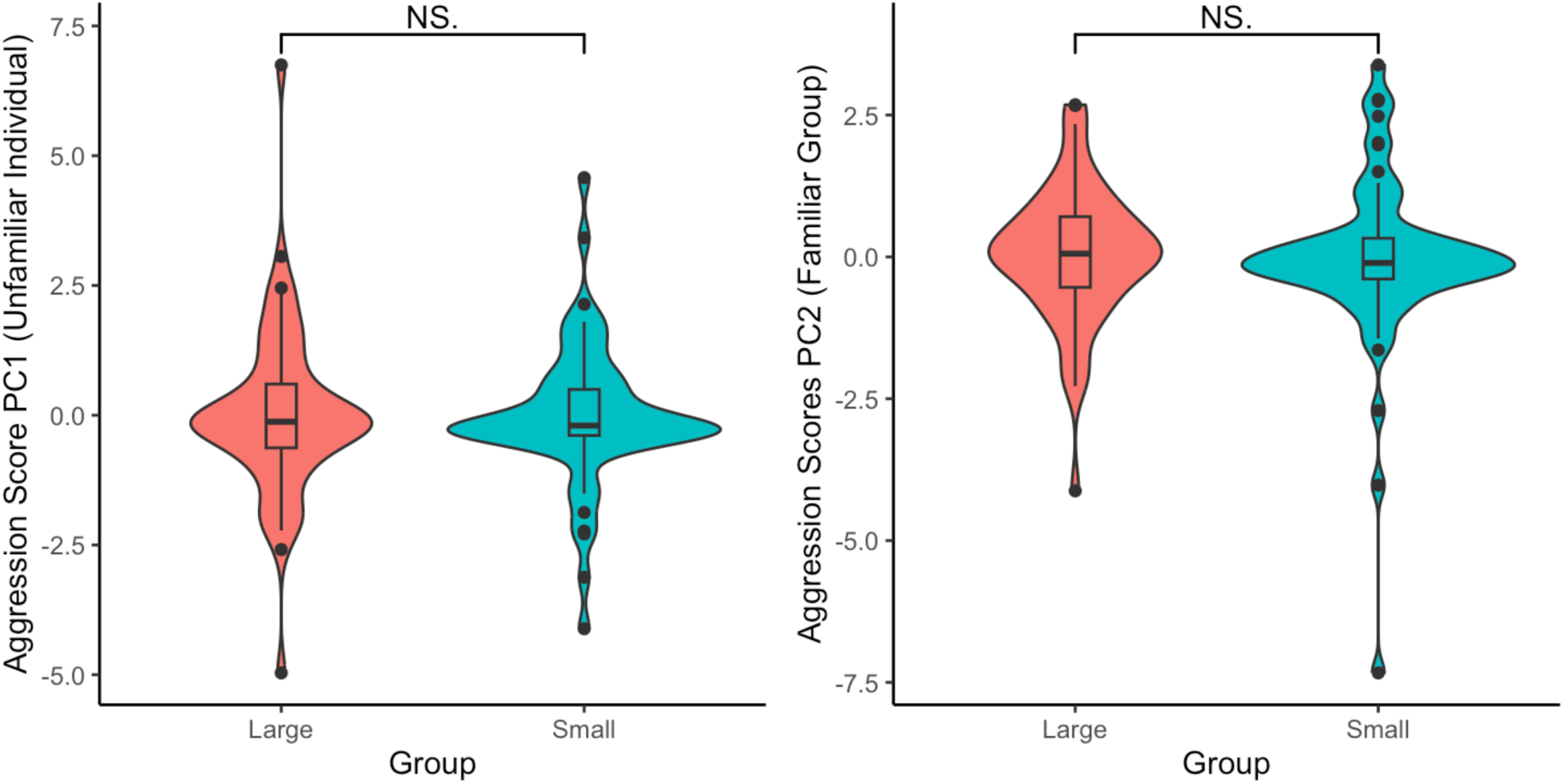
There were no significant differences in aggression scores between social groups a) during the Unfamiliar Individual (*χ^2^* = 0.001, *d.f.* = 1, *p* = 0.973), and b) during the Familiar Group task (*χ^2^* = 0.183, *d.f.* = 1, *p* = 0.670). *Note: for the axis PC2, more aggressive individuals have a negative score.

Aggression scores for the Familiar Group task (PC2) were not explained by Group Size (*χ^2^* = 0.183, *d.f.* = 1, *p* = 0.670; Fig. 6). However, males were significantly more aggressive than females (Females: 0.35 ± 0.97, Males: -0.35 ± 1.61; *χ^2^* = 8.554, *d.f.* = 1, *p* = 0.004; Figure S5), and quails with a higher weight were significantly more aggressive than quails with a lower weight (*χ^2^* = 6.002, *d.f.* = 1, *p* = 0.016; Figure S6).

## 5. Discussion

The main goals of this study were threefold: (A) to examine the link between response inhibition and impulsive aggression at an individual level, and (B) to assess whether group size during early-life environment influenced the development of response inhibition and (C) impulsive aggression using juvenile Japanese quails. We found (A) some significant correlations between measures of response inhibition and impulsive aggression, providing some limited support to the theoretical model linking both concepts. We also found (B) evidence that early-life social environment indeed shapes the development of response inhibition, as quails raised in smaller social groups spent more time repeating an inefficient action (i.e., pecking at the cylinder) than those raised in larger social groups. However, contrary to what we expected, (C) we found no evidence that early-life social environment influenced the development of impulsive aggression.

### Relationship between response inhibition and impulsive aggression

In juvenile Japanese quails, we found some evidence that response inhibition was significantly linked to impulsive aggression at an individual level. More specifically, increased perseveration on an unrewarded action towards an unfamiliar object (i.e., time spent pecking at the transparent cylinder) was significantly correlated to increased aggression towards unfamiliar individuals. This result may suggest that individuals tend to interact aggressively towards novelty, be it an object or an individual. The link between higher aggression and lower inhibition has also been found in previous observational studies in mammals (12,13,71,72), as suggested by theoretical models (14). In primates, aggressive individuals with low response inhibition are thought to have an advantage to access and retain food resources (73). A similar observation was found in Australian magpies, where individuals involved in aggressive interactions (both initiating and receiving) performed worse in an associative learning task, possibly due to lower time and/or energy to attend the task (74). While the authors did not test inhibition directly, previous work in that study population has found a significant correlation between inhibition (as assessed with a cylinder) and associative learning (31). Together, these results highlight how the social environment, its dynamics, and cognitive performance are tightly linked with one another.

Interestingly, we also found that individuals that successfully fully cancelled a discrete action (i.e., by detouring without pecking at the transparent barrier) had significantly increased levels of aggression towards familiar individuals. This result goes in the opposite direction to what we predicted, and contrast with previous studies that investigated the link between inhibition and aggression. One potential explanation for this seemingly contradictory result could be by viewing the adaptive significance of aggression (3). Indeed, aggressive individuals are more likely to initiate contests and to subsequently win them (75), which potentially gives them higher fitness payoffs. Hence, individuals with better response inhibition may use the most advantageous coping strategy in response to the social environment demands.

These contradictory results may suggest three things: a) that the mechanisms involved during the cancellation of an action depend on whether the individuals are already engaged in the action they need to inhibit, and/or b) that the tasks are not assessing response inhibition or aggression, but rather pecking tendency (whether it is towards an object or an individual), and/or c) that response inhibition and aggression depends highly on the social context.

Indeed, when examining in detail the requirements of both the Detour and the Cylinder tasks, individuals may have to inhibit two actions: first, the prepotent response of going directly at the food, thus pecking at the transparent barrier, and second, the repetitive action of pecking along the barrier if they failed to inhibit the first action (76). The lack of correlations between the different measures of the two response inhibition tasks occurred despite both tasks requiring individuals to detour around a transparent barrier (47) and despite both tasks having a similar “go” stimulus (i.e., the visible food reward) and a similar “stop” stimulus (i.e., the transparent barrier) present at the same time (76). Rather, the characteristics of the stop stimulus itself, such as the shape (e.g., (77)) and the motor action required to successfully detour, might influence the individual’s response: the transparent barrier in the Barrier task is large and requires individuals to move their entire body around it, whereas the transparent barrier in the Cylinder task is small and requires individuals to place their head through the openings (55). Even when carefully choosing tasks purporting to assess the same aspect of response inhibition, our results suggest that response inhibition is multi-faceted and context-dependent (55,78–80). Not only the nature of the action to be stopped, but the overall task context in which actions need to be stopped could thus involve different mechanisms.

Alternatively, training differences and learning may explain the lack of repeatability in performance we observed. For the Barrier task, individuals were trained to go around the opaque barrier and might have learned the detouring solution (81), explaining the high success of individuals in this task during the first transparent trial. In comparison, quails were not trained with an opaque cylinder during the Cylinder task. Hence, quails had to not only inhibit pecking at the transparent cylinder, but also to successfully “solve” the task (i.e., learning the detouring solution (55,56,82). This may also explain why the number of quails who successfully detoured during the Cylinder task was lower than during the Barrier task. Thus, the cognitive processes involved during both tasks may have differed. Additionally, unlike what was found with pheasants (78), there was no evidence that Japanese quails used their previous experience of detouring around a transparent barrier, as detouring success and time spent pecking during the Cylinder task did not improve from during the Barrier task, supporting once more the idea that response inhibition is context-dependent. Understanding how and why exactly there is such a low reliability between tasks is beyond the scope of the current study but warrants further investigation (83).

Similarly, the contradictory results we obtained regarding the link between aggression and response inhibition may be due to the contextual differences between both aggression tasks. Individuals’ behaviour may differ when they are in the presence of familiar and of unfamiliar individuals (e.g., (84), “winner and loser effects” – (85), “dear enemy” hypothesis – as reviewed by (86); but see (44) for a counter-example with male Japanese quails). Because agonistic interactions between individuals can be costly, familiarity is thought to play a key role in the formation of hierarchies, both within species (87) and between species (88). For this reason, aggression is usually reduced within the group once hierarchies are established (3). Yet, the lack of repeatability in individual aggression scores is surprising as it goes against the idea of aggression as a personality trait (e.g., (89–91)). Instead, this result suggests that aggression is subject to high intra-individual variation (92). Plasticity in aggressive responses could depend on the outcome of past antagonistic interactions (93) and on the traits of the other individual, such as status signals or signals of fighting ability (reviewed by (94)) or use social information to assess the costs of engaging in an antagonistic behaviour towards another individual (95).

Regardless of the lack of correlations in aggressive behaviours between tasks, we found that behaviours associated with individuals that were the target of aggressive behaviours (as assessed by the time spent escaping and the number of pecks received) were less likely to engage in aggressive behaviours (as assessed by the time spent chasing and pushing, and the number of pecks given) within each task context. While we did not assess hierarchical dominance directly, this behavioral trend suggests differences in social rank between individuals. Hierarchical dominance has been found to influence, or even suppress, the expression of certain cognitive abilities and to affect brain morphology ((96–99), but see (100)), and may contribute to the duality in our results linking response inhibition and aggression.

The social context of the task may also explain this duality in our results. Aggression was assessed in a group context whereas response inhibition was assessed in a solitary context. The social context may exert different inhibitory pressures on individuals. For instance, in the presence of a more dominant or more aggressive group member, an individual might inhibit going towards the food reward to reduce the risk of receiving directed aggression (101). However, when alone, that same individual may or may not have lower response inhibition levels towards the food rewards. This possibility suggests, once again, the importance of accounting for the social context when evaluating impulsivity generally.

### Influence of early-life social environment on response inhibition and impulsive aggression

The importance of the social environment on the development of cognition has been suggested in many species (20,21,23,30,74,102). More specifically, social group size, a key component of the social environment, has been found to correlate with higher levels of inhibition in hyenas (32) and in Australian magpies (31), but not in guppies (34,103) nor in horses (104). Here, we found that quails raised in larger groups inhibited a repetitive unrewarded action (i.e., pecking at the transparent cylinder) faster, on average, than quails raised in smaller groups. This result aligns with those from previous studies evaluating intraspecific variability in response inhibition using the Cylinder task on Australian magpies and on hyenas (31,32), and support what we predicted. However, we found no significant effect of social group size on the overall success rate for both tasks, nor on the time spent pecking on the transparent barrier during the Barrier task. As we previously mentioned, both tasks may not measure the same aspect of response inhibition or may measure pecking tendency in one or the other. Therefore, the influence of the social environment might be mediated differently depending on which aspect of response inhibition is considered.

Early-life social environmental conditions have strong effects on aggressive behaviour in many species ((42,105–107)), with usually a decrease in aggression and higher social tolerance as group size increases (e.g., (37,40,49)). Unlike previous research, we found no significant difference in levels of aggression between groups during our aggression tasks. This lack of significant results might be caused by the increase in density associated with the increase in group size, as individuals of both conditions were given the same size enclosure. Higher population density has been associated with higher aggression levels (e.g., lizards – (108)), as individuals switch to a resource defence strategy (36,37). Yet, unlike group-living species in the wild that may encounter limited amount of food, our quails were provided with *ad lib* access to food (outside of food restriction procedures). Resource availability and competition for food should have been the same for both groups, independently of group density and size. In such circumstances, theoretical models suggest that aggression levels should be low for both groups (109), and that individuals should be socially tolerant (i.e., tolerance hypothesis; (110)). Disentangling the effects of group size and group density on aggression as well as on response inhibition is a promising next step.

Social group size is not the only aspect of social complexity. Group dynamics and stability, as well as the nature of the relationships between group members, are other important aspects that contribute to the variation in social complexity within and between groups (29,50). Further investigation is needed to determine which aspects of the social life influence the development of aggression and of response inhibition (111).

### Intra- and inter-individual variation in response inhibition and in impulsive aggression

Research on non-human animals had, until recently, tried to minimize the importance of inter-individual variation in cognition (112,113). Now, there is a general effort in pinpointing which factors may underlie individual differences in cognition generally (114,115). Despite controlling the early-life physical and social environment of our quails, as well as the acclimatation, habituation and testing procedures, we still observed pronounced individual differences in the success and time spent performing a repetitive unrewarded action during the response inhibition tasks and in the aggressive behaviours during the impulsive aggression tasks that were not explained by the difference in group size.

These inter-individual differences in response inhibition and impulsive aggression may be due to differences in motivation and attention (116–118). Some individuals might have greater motivation to participate during the tasks due and interact with apparatuses and/or other individuals due to high exploration/low neophobia (119–123) or due to higher basal energetic demands than others (124). The latter could explain why we found an effect of body weight on time spent pecking at the transparent barrier (125). However, given that the direction of that effect was opposite for both response inhibition tasks, we cannot confirm this explanation as to why or how it might be the case. Additionally, in group-living domestic species, more aggressive individuals often outcompete less aggressive individuals for limited resources and have a higher weight as a result (e.g., (126)), which support our findings on heavier individuals being more aggressive towards familiar individuals. Concurrently, being aggressive might be less costly and more beneficial for larger individuals, as they are more likely to physically outcompete smaller ones.

Cognitive abilities and personality traits have also been known to differ between females and males in some species (127–131). Despite not finding any differences in response inhibition between males and females, we found that males were significantly more aggressive than females (44,132–136). We found such a sex difference in aggression when the quails were between 5- and 6-weeks old, at the onset of sexual maturity in Japanese quails (between 6- to 8-weeks old). Little is known regarding aggression in immature, non-sexually active or castrated quails, but it is usually assumed they have lower aggressive tendencies than sexually mature ones (43,137). Hence, inter-individual variation in aggression that we observed might also be due to inter-individual differences in reaching sexual maturity, despite keeping all other parameters constant for all individuals (photoperiod, temperature, food). Similarly, in our study, response inhibition was assessed in juvenile quails. Previous research on other species show that inhibition follows a developmental period (138,139). While there has not been a study directly investigating the development of inhibition in Japanese quails, domestic chicks (*Gallus gallus*), a closely related precocious species who follows a similar developmental pattern (both species reaching maturity at six to eight weeks old), can already detour when they are two weeks old (138). As quails were at least three weeks old at the time of testing, we expected them to have fully completed the developmental period in response inhibition, but that assumption might be erroneous. Hence, the lack of consistent correlations we found between measures in response inhibition and aggression may be due to one or both traits not being fully developed when assessed.

Finally, a large inter-individual variation in inhibiting a behavioral response might indicate a lack of strong selection on a given trait (e.g.,(140)), or alternatively, that it might be favored in certain environments or contexts, but not in others (141,142). Similar arguments can be made to explain intra-individual variation in response inhibition and in aggression. Indeed, the lack of repeatability in response inhibition at an individual level may suggest that individuals are flexible in their cognitive ability and in their social behavior, depending on the context. For instance, impulsive actions might be more adaptive in an unstable or unpredictable environment or when travel time between resources increase (143). Understanding in which contexts extreme values of response inhibition might be favored and why such variation in cognitive traits and in personality exists within the population is a topic that warrants further investigation.

## 6. Conclusion

In summary, our study provides one of the first experimental evidence that early-life social environment can influence the development of response inhibition. We also experimentally showed that certain aspects of aggression are linked to deficits in response inhibition, especially regarding perseverating in performing an unrewarded action towards novelty (objects and/or individual). Hence, our study shed some light regarding the mechanisms underlying impulsive aggression using Japanese quails. Yet, contrary to what was expected based on previous literature, group size, a key aspect of early-life social environment, did not influence aggression in our quails, despite it influencing certain aspects of response inhibition (the same ones that were linked to aggression). Aggression is a multifaceted and complex behaviour that involves multiple processes (144,145). Thus, despite controlling the physical and social environment, intrinsic factors such as genetics, motivation, non-social personality traits (i.e., neophobia, exploration, activity), and the ability to get social information might explain inter-individual variability in aggression that we witnessed and mediate the link between response inhibition and aggression. Regarding the influence of sociality on the emergence of aggressive behaviors, other social factors, such as group dynamics and stability, and further research is warranted to fully understand how sociality, response inhibition, and aggression relate to one another.

## Acknowledgements

We thank the staff of the Wildlife Rescue Centre Ostend for general project support and Harry Suter for insightful discussions and comments on the draft of the manuscript. We also thank two anonymous reviewers for their critical and insightful comments.

## Ethical Statement

The experiment was performed in accordance with the Association for the Study of Animal Behaviour ethical guidelines under permission of the ethical committee of animal experimentation (VIB Site Ghent, Universiteit Gent): EC2022-005.

## Data Accessibility

All videos of the experiment can be found on Zenodo (https://zenodo.org/records/13361318).

R codes and extracted data from BORIS can be found on OSF (https://osf.io/2b5uy/).

## Author contributions

Based on the CRediT system, the contributions of the authors are as follow:

‘Conceptualization’ – AV, LL, FV; ‘Data curation’ – AV, KW; ‘Formal analysis’ – AV; ‘Funding acquisition’ – AV, AD, WZ, CAT, AM, LL, FV; ‘Investigation’ – AV, KW, RA, AD, WZ, CAT, SK, LL, FV, ‘Methodology’ – AV, KW, RA, AD, LL, FV; ‘Project administration’ – AV, FV; ‘Resources’ – FV, ‘Supervision’ – AV, LL, FV; ‘Validation’ – AV, KW, SK, CAT; ‘Visualization’ – AV, ‘Writing—original draft’ – AV, and ‘Writing—review & editing’ – AV, KW, RA, AD, WZ, CAT, AM, LL, FV. In the Methods section, we followed the MERIT system (Nakagawa et al. 2023).

