## Supplementary Material for "To Peck or Not To Peck: The influence of early-life social environment on response inhibition and impulsive aggression in Japanese quails"

Alizée Vernouillet ORCID: <https://orcid.org/0000-0003-2525-2936>

Kathryn Willcox ORCID: <https://orcid.org/0000-0002-3222-2850>

Reinoud Allaert ORCID: <https://orcid.org/0000-0002-6122-496X>

Anneleen Dewulf ORCID: <https://orcid.org/0000-0003-4215-0421>

Wen Zhang ORCID: <https://orcid.org/0000-0001-9120-9763>

Camille A. Troisi ORCID: <https://orcid.org/0000-0002-4036-3848>

Sophia Knoch ORCID: <https://orcid.org/0000-0001-9156-5725>

An Martel ORCID: <https://orcid.org/0000-0001-7609-5649>

Luc Lens ORCID: <https://orcid.org/0000-0002-0241-2215>

Frederick Verbruggen ORCID: <https://orcid.org/0000-0002-7958-0719>

**Supplementary methods**

### *General Experimental Procedures*

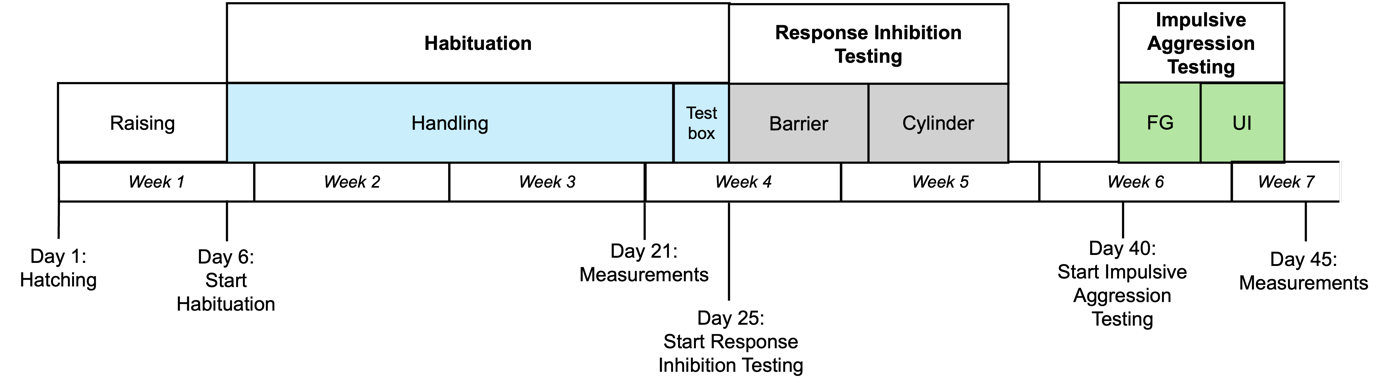

Figure S1: Experimental timeline of the study with 120 Japanese quails. All quails hatched on the same day. FG – Familiar Group task, UI – Unfamiliar Individual task.

### *Impulsive Aggression tasks*

Table S1: Ethogram of the different behaviors coded during the Response Inhibition tasks and the Impulsive Aggression tasks in Japanese quails.

| **Tasks** | **Behavior** | **Description** |
| --- | --- | --- |
| *All* | Leaving start box | The bird’s whole body is out of the start box |
|  | Eating | The bird’s head enters the reward feeder or the bird pecks at the worm |
| *Response Inhibition* | Detour | The bird’s front half of the body has passed the imaginary line in the continuation of the barrier (Barrier task)  The bird’s head entered the side opening of the cylinder (Cylinder task) |
| *Impulsive Aggression* | Pecking | The bird uses its beak in a pecking manner, raising its head and vigorously touching the other bird |
|  | Chasing | The bird is moving at a greater speed than walking towards another bird and following it |
|  | Escaping | The bird is moving at a greater speed than walking, such that it moves away from another bird |

**Supplementary statistical analyses**

*Structural Equation Modeling*

We wanted to examine the relationship between variables using a SEM with the structural equation approach (schematic of the model represented in Fig. S2) using the package *lavaan* (Rosseel 2012). The following observed variables were used to construct structural equation models (SEMs): the time spent ‘interacting’ with the barrier/cylinder (i.e., time the bird spends actively and repeatedly pecking/pushing the barrier) during each Response Inhibition task and the number of pecks given, the number of pecks received, the time spent chasing, the time spent escaping, and the time spent pushing during each Impulsive Aggression task. Performance during each Response Inhibition task was not included, as this measure was binary (success/fail). We also included morphology measures as part of a Morphology factor that could influence Response Inhibition and Impulsive Aggression, and in turn that could be influenced by Group Size. However, before analysing the data using an SEM, we performed a Confirmatory Factor Analysis to determine whether the observed variables assessed the latent factors of interest (Response Inhibition, Impulsive Aggression, Morphology).

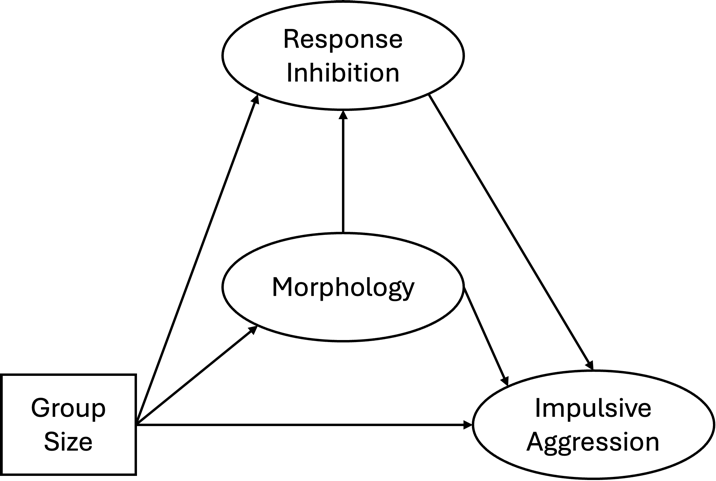

Figure S2: Planned schematic of the relationships estimated by the Structural Equation Modelling.

*Factor analyses (FA)*

A first Confirmatory Factor Analysis was done using the package *psych* (Revelle 2024). We tested a first hypothesized grouping of factors based on the current literature and the model included three latent factors (Fig. S3).

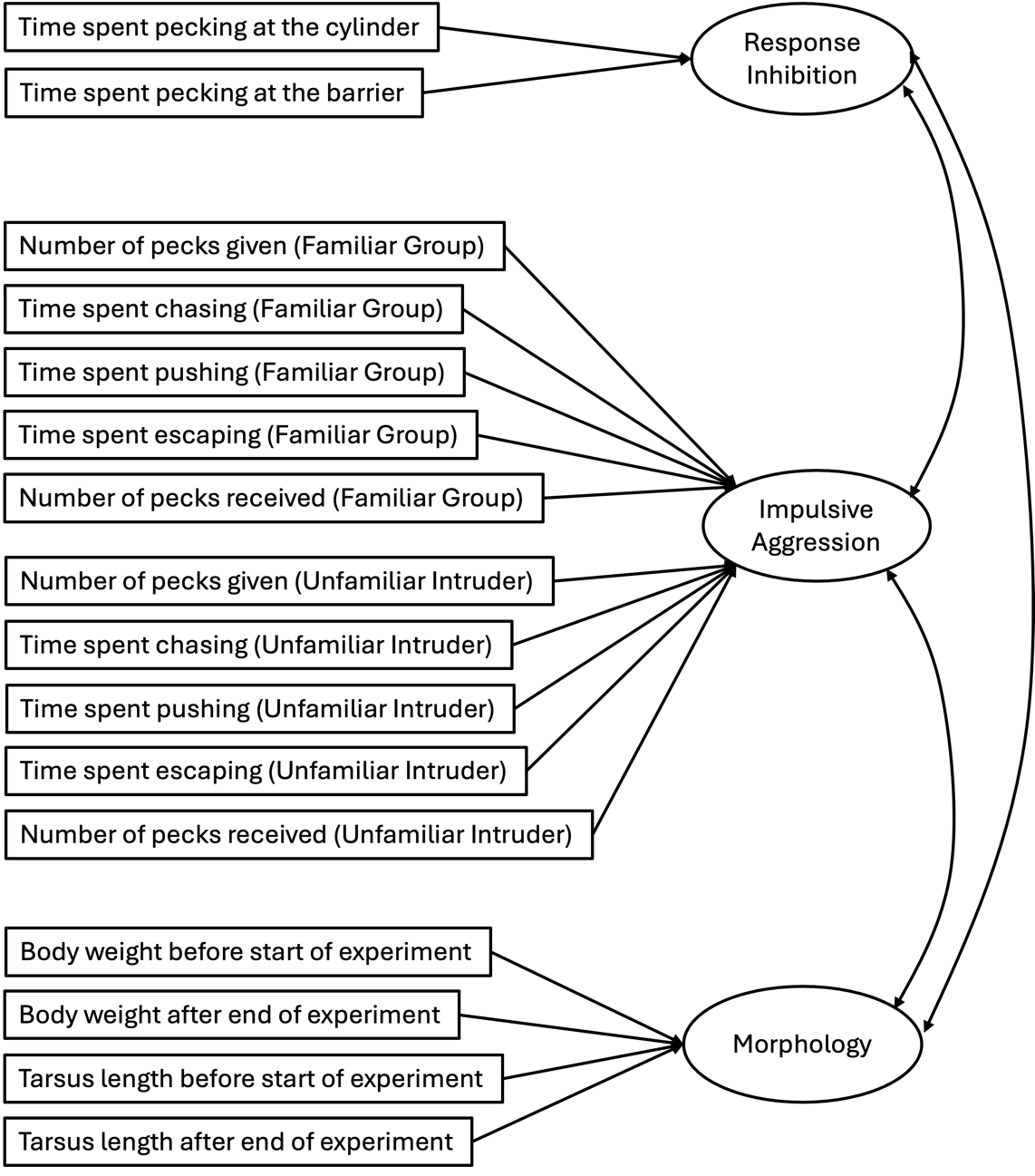

Figure S3: Initial hypothesized model tested using the Confirmatory Factor Analysis. Observed variables are in rectangles and latent variables of interest are in circles.

The following observed variables were used during our analyses: the time spent ‘interacting’ with the barrier/cylinder (i.e., time the bird spends actively and repeatedly pecking/pushing the barrier) during each Response Inhibition task for the Response Inhibition factor, the weight and the length of the right tarsus taken before and after the experiment for each bird for the Morphology factor, and the number of pecks given, the number of pecks received, the time spent chasing, the time spent escaping, and the time spent pushing during each Impulsive Aggression task for the Impulsive Aggression factor (see Fig. S3). Performance during each Response Inhibition task was not included, as this measure was binary (success/fail), and factor analyses can only be performed on continuous variables.

Given that factor analyses are sensitive to outliers, we examined the distribution of each variable and removed obvious outliers (i.e., when the histogram showed a clear break in the distribution). As a result, we removed data from 14 birds for this analysis. Additionally, factor analyses cannot be performed on missing values, therefore we removed an additional 22 birds from this analysis (i.e., individuals that did not participate at either Response Inhibition task). Hence, the factor analyses used a reduced dataset of 84 birds.

We used the following criteria to evaluate the models fit: Comparative Fit Index (*CFI*) > 0.95, Tucker-Lewis Index (*TLI*) > 0.95, the root mean square error of approximation (*RMSEA*) < 0.06, the standardised root mean square residual (*SRMR*) < 0.08, and *chi-square*.

CFA results from that first model indicated a clear lack of fit, with the following fit indices: *CFI* = 0.679, *TLI* = 0.618, *RMSEA* = 0.128, *SRMR* = 0.125, and *chi-square* = 239.06, *df* = 101, *p* = 0.000. The Morphology factor was well defined, but the Impulsive Aggression and the Response Inhibition factors were not.

We then used an Exploratory Factor Analysis (EFA) approach to determine how many factors should be defined in our dataset and which variables should be grouped in each factor. The model identified that four factors would be the optimal number of factors to explain 51.9% of the variation of the data (*chi-square* = 101.11, *df* = 62, *p* = 0.001). Factor extraction results can be found in Table S2. Based on the loadings of each variable for each factor, we can identify a Morphology factor (Factor 1) with all morphology factors, an Impulsive Aggression towards unfamiliar individuals factor (Factor 2) with the number of pecks given and the time spent chasing during the Unfamiliar Individual fast, an Impulsive Aggression towards familiar individuals factor (Factor 3) with the number of pecks given and the time spent pushing during the Familiar Group task, and an Aggression Received/Submission from familiar individuals factor (Factor 4) with the number of pecks received and the time spent escaping during the Familiar Group task, as well as the time spent pecking during the Cylinder task.

Table S2. Factors extraction during the Exploratory Factor Analysis. FG = Familiar Group task, UI = Unfamiliar Individual task. In bold are variables whose loadings are higher that 0.4 (hence, highlighting significant loadings).

|  | Factor 1 | Factor 2 | Factor 3 | Factor 4 |
| --- | --- | --- | --- | --- |
| Number of pecks given during FG |  | -0.120 | **0.893** | -0.248 |
| Number of pecks received during FG |  | -0.162 |  | **0.510** |
| Time spent chasing during FG |  |  | 0.374 |  |
| Time spent escaping during FG | -0.148 | -0.161 | -0.322 | **0.813** |
| Time spent pushing during FG |  | -0.137 | **0.679** | -0.177 |
| Number of pecks given during UI |  | **0.757** | 0.140 |  |
| Number of pecks received during UI |  | -0.144 | -0.167 | -0.259 |
| Time spent chasing during UI |  | **1.079** | -0.241 | -0.174 |
| Time spent escaping during UI |  | -0.143 | -0.228 | -0.318 |
| Time spent pushing during UI |  | -0.139 | 0.278 | 0.184 |
| Time spent pecking during the Cylinder task | 0.160 | 0.110 |  | **0.434** |
| Time spent pecking during the Barrier task | -0.343 |  | -0.173 |  |
| Tarsus length measured at start of the experiment | **0.909** |  |  | -0.178 |
| Weight measured at start of the experiment | **0.916** | -0.122 | 0.122 |  |
| Weight measured at end of the experiment | **0.796** |  | -0.166 |  |
| Tarsus length measured at end of the experiment | **0.741** |  | -0.204 |  |

Based on these results, we refined our model for the Confirmatory Factor Analysis by making it more context specific. We tested a second model that included the latent factors Response Inhibition, Impulsive Aggression during the Familiar Group task, Impulsive Aggression during the Unfamiliar Individual task, Aggression Received during the Familiar Group task, Aggression Received during the Unfamiliar Individual task, and Morphology. The number of pecks received and the time spent escaping were used as observed measures of the Aggression Received factor and the number of pecks given, the time spent chasing and the time spent pushing were used as observed measures of the Impulsive Aggression factor for each task. While the second model indicated a clear improvement in comparison to the first model using Aikaike indices (First model AIC: 7904.467, Second model AIC: 7826.499), the second model still showed a lack of fit, as seen with the following fit indices: *CFI* = 0.888, *TLI* = 0.849, *RMSEA* = 0.080, *SRMR* = 0.097, and *chi-square* = 137.088, *df* = 89, *p* = 0.001.

Given the lack of support for different groupings, we did not continue forward with our planned SEM analyses. We instead used a correlation approach and a Principal Component Analysis (PCA) approach to investigate the contextual consistency of Response Inhibition and Impulsive Aggression respectively. We then assessed the relationships between Response Inhibition and Impulsive Aggression using a correlation and a t-test approach. Finally, we assessed the influence of Group Size, morphology, and sex on different measures of Response Inhibition and scores of Impulsive Aggression using once again a GLM approach. More detailed explanations are included in the main text.

**Supplementary Results**

Table S3. Overview of the quails based on whether they detoured and/or pecked on the transparent barrier or cylinder during the Barrier task and the Cylinder task, depending on their sex and the social group they were raised in (*n* = 120).

|  |  | Sex (F/M) | Group (Small/Large) | Total |
| --- | --- | --- | --- | --- |
| Barrier task | Detoured without pecking | 6/13 | 11/8 | 19 |
|  | Detoured but pecked | 51/42 | 43/50 | 93 |
|  | Did not detour but pecked | 2/5 | 5/2 | 7 |
|  | Did not detour nor pecked | 1/0 | 1/0 | 1 |
| Cylinder task | Detoured without pecking | 8/6 | 5/9 | 14 |
|  | Detoured but pecked | 24/22 | 19/27 | 46 |
|  | Did not detour but pecked | 19/17 | 21/15 | 36 |
|  | Did not detour nor pecked | 9/15 | 15/9 | 24 |

**Supplementary figures**

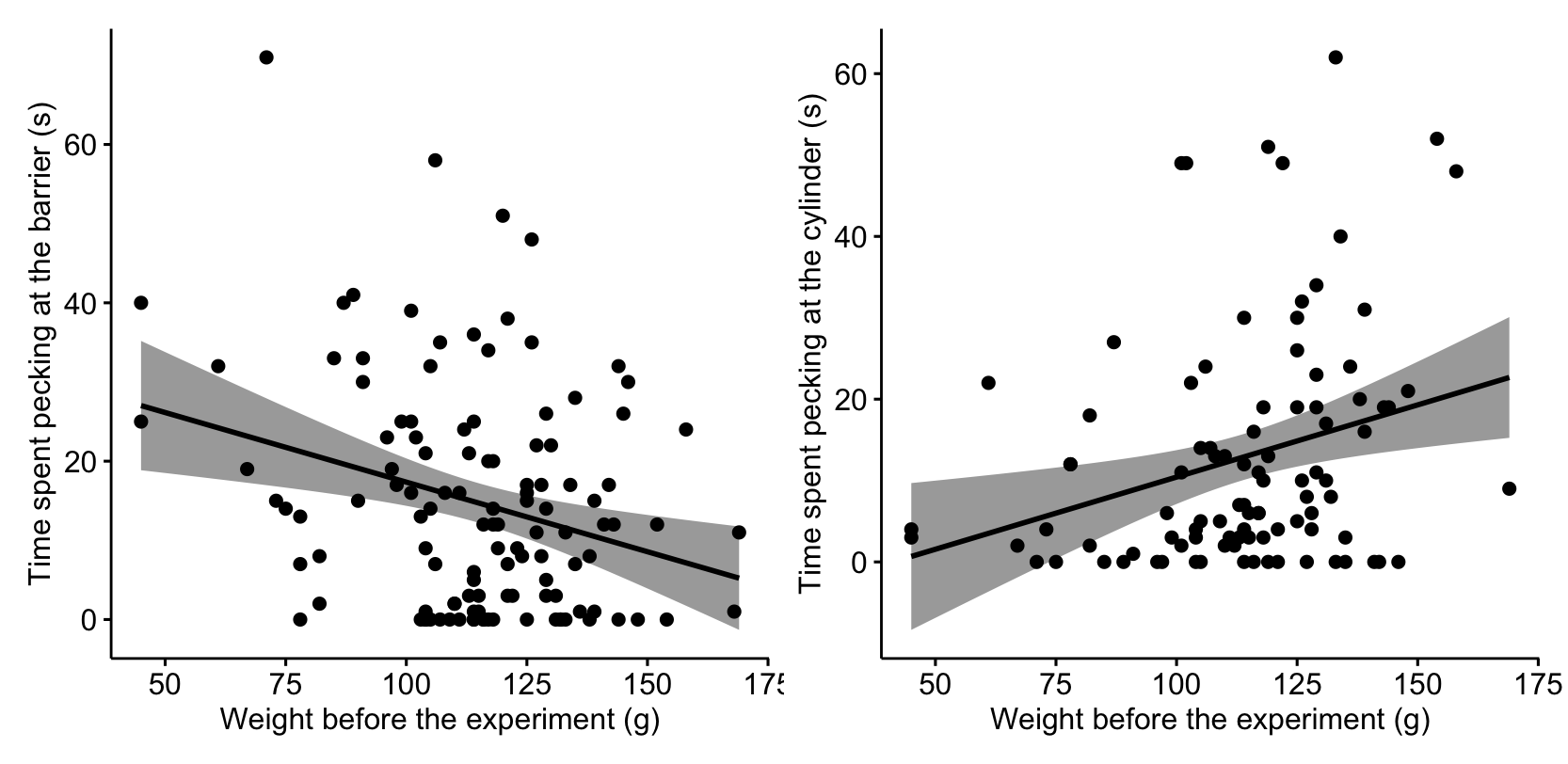

Figure S4. Quails with a lower body weight spent more time pecking at the transparent barrier during the Barrier task (*χ^2^* = 5.817, *d.f.* = 1, *p* = 0.016, left) and significantly less time pecking at the transparent cylinder during the Cylinder task (*χ^2^* = 13.078, *d.f.* = 1, *p* < 0.001, right) than quails with a higher body weight.

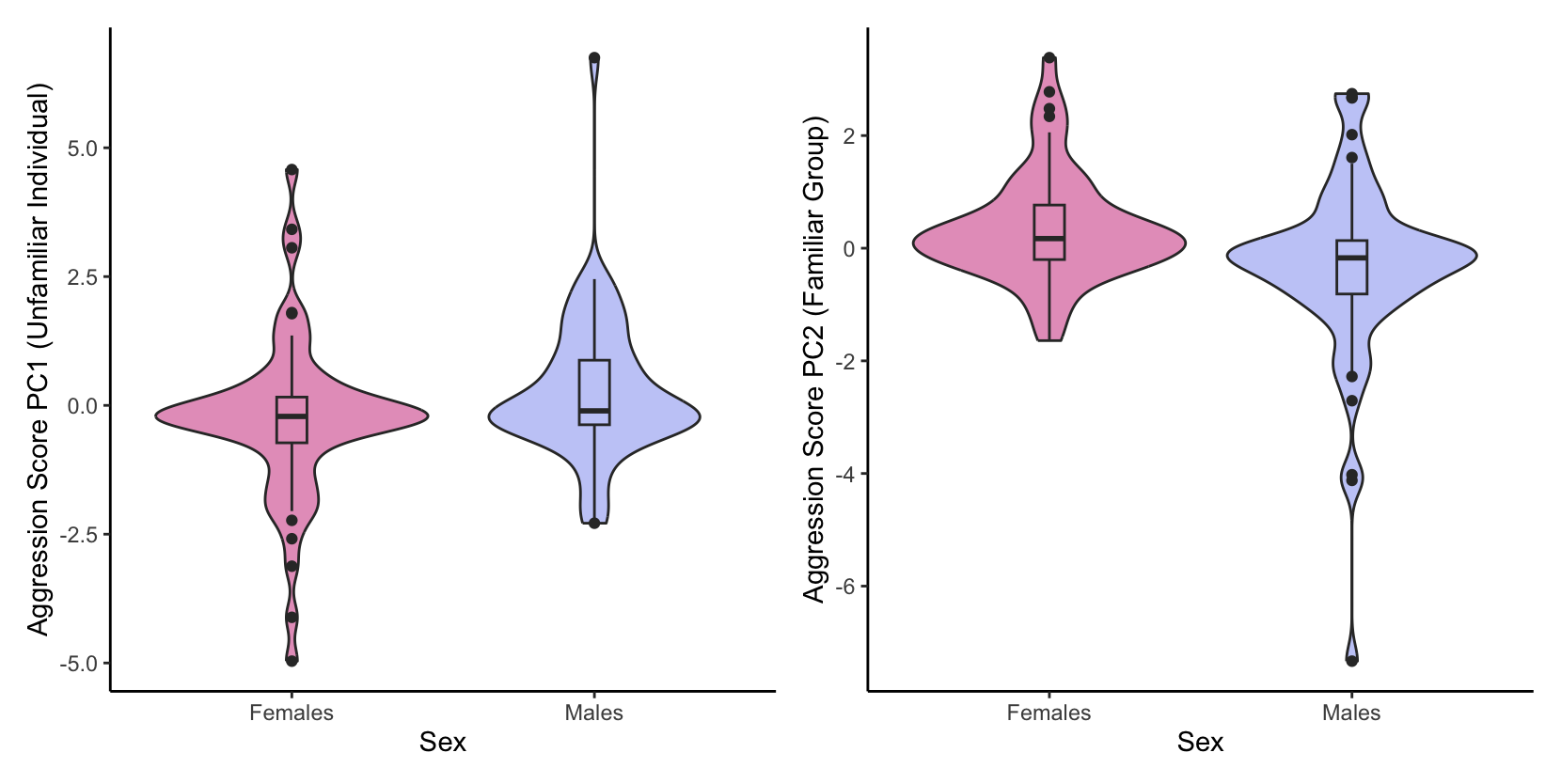

Figure S5. Male Japanese quails have a significantly higher aggression score a) during the Unfamiliar Individual (*χ^2^* = 4.318, *d.f.* = 1, *p* = 0.038), and b) during the Familiar Group task (*χ^2^* = 8.554, *d.f.* = 1, *p* = 0.004) than females. *Note: for the axis PC2, more aggressive individuals have a negative score.

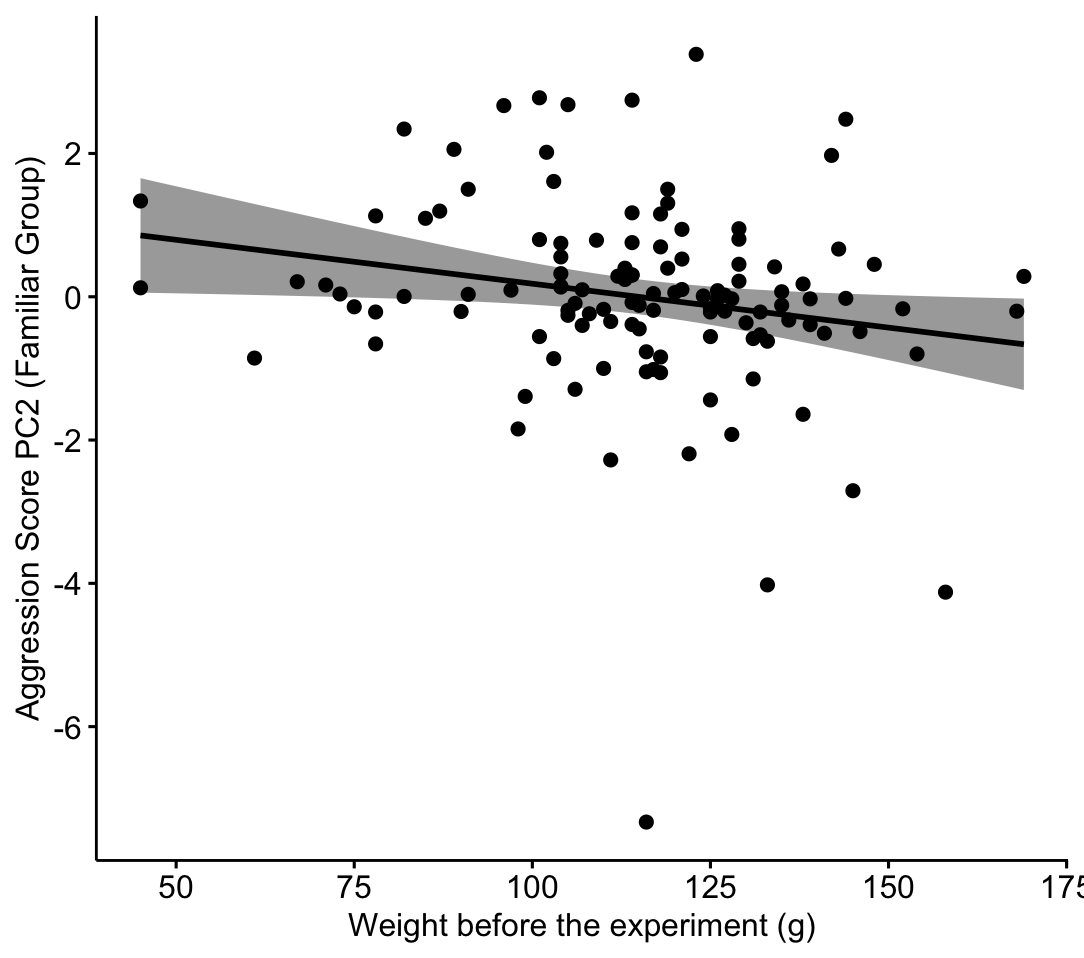

Figure S6. Quails with a higher body weight were significantly more aggressive during the Familiar Group (*χ^2^* = 6.002, *d.f.* = 1, *p* = 0.016).

Rosseel Y (2012). “lavaan: An R Package for Structural Equation Modeling.” *Journal of Statistical Software*, **48**(2), 1–36. [doi:10.18637/jss.v048.i02](https://doi.org/10.18637/jss.v048.i02).

Revelle W (2024). *psych: Procedures for Psychological, Psychometric, and Personality Research*. Northwestern University, Evanston, Illinois. R package version 2.4.6, [https://CRAN.R-project.org/package=psych](https://cran.r-project.org/package=psych).
